# The erythrocyte membrane properties of beta thalassaemia heterozygotes and their consequences for *Plasmodium falciparum* invasion

**DOI:** 10.1101/2022.01.13.476158

**Authors:** Viola Introini, Alejandro Marin-Menendez, Guilherme Nettesheim, Yen-Chun Lin, Silvia N. Kariuki, Adrian L Smith, Letitia Jean, John N. Brewin, David C. Rees, Pietro Cicuta, Julian C. Rayner, Bridget S. Penman

## Abstract

Malaria parasites such as *Plasmodium falciparum* have exerted formidable selective pressures on the human genome. Of the human genetic variants associated with malaria protection, beta thalassaemia (a haemoglobinopathy) was the earliest to be associated with malaria prevalence. However, the malaria protective properties of beta thalassaemic erythrocytes remain unclear. Here we studied the mechanics and surface protein expression of beta thalassaemia heterozygous erythrocytes, measured their susceptibility to *P. falciparum* invasion, and calculated the energy required for merozoites to invade them. We found invasion-relevant differences in beta thalassaemic cells versus matched controls, specifically: elevated membrane tension, reduced bending modulus, and higher levels of expression of the major invasion receptor basigin. However, these differences acted in opposition to each other with respect to their likely impact on invasion, and overall we did not observe beta thalassaemic cells to have lower *P. falciparum* invasion efficiency for any of the strains tested.

## Introduction

Selection for resistance to malaria disease has elevated the frequencies of a wide variety of human genetic variants, most famously genetic disorders of haemoglobin^1^. Haldane was the first to suggest that high frequencies of a haemoglobinopathy, beta thalassaemia, could reflect a history of malaria exposure in certain populations^2^. In 1954 Anthony Allison confirmed Haldane’s hypothesis for another haemoglobinopathy: sickle cell anaemia^3^. A large body of evidence has since confirmed the profound protective effects of sickle cell heterozygosity against both uncomplicated and severe *Plasmodium falciparum* malaria, as well as protective effects of two additional haemoglobinopathies (alpha thalassaemia and haemoglobin C) against severe *P. falciparum* malaria^4^. However, despite Haldane’s malaria hypothesis originally referring specifically to beta thalassaemia, far less evidence is available to confirm beta thalassaemia’s malaria protective effect. This is because the genetics of malaria resistance have been most intensely studied in sub-Saharan Africa, where beta thalassaemia is relatively uncommon. The main evidence for the malaria protective effect of beta thalassaemia remains its geographical co-location with the historical incidence of *P. falciparum* malaria in Southern Europe, North and West Africa and Asia. In Sardinian villages, beta thalassaemia frequencies inversely correlate with altitude and hence historical malaria exposure^5^. A case control study conducted in Liberia also suggests that heterozygosity for beta thalassaemic mutations offers protection against hospital admission with malaria^6^. Beta thalassaemia is caused by mutations which reduce or eliminate beta globin production from the beta globin locus (*HBB*). The pathophysiology of beta thalassaemia results from an excess of unpaired alpha globin chains accumulating in the erythrocytes of affected individuals and causing oxidative damage^7^. Heterozygosity for beta thalassaemic mutations, known also as beta thalassaemic minor or beta thalassaemia trait, results only in a mild anaemia, but homozygosity or compound heterozygosity can cause thalassaemia major, a life threatening transfusion dependent condition.

Beta thalassaemia heterozygotes typically have erythrocytes with smaller mean corpuscular volume (MCV) and mean corpuscular haemoglobin than wild type cells^8^. Known alterations to the membrane of red blood cells in beta thalassaemia heterozygotes include an increase in the flux of K^+^ across the membrane^9^, which is also observed in iron deficiency anaemia and sickle cell disease. There likewise may be altered lipid content in heterozygous beta thalassaemic erythrocyte membranes, similarly mirroring changes seen in iron deficiency anaemia^10^.

In severe (homozygous or compound heterozygous) beta thalassaemia, erythrocyte membranes are mechanically unstable (where stability measures the resistance of membranes to fragmentation^11^), and Schrier and Mohandas^12,13^ have shown that this could result from the association of oxidised unpaired alpha globin chains with the erythrocyte membrane. The extent to which this impacts the biophysical properties of less-severely-affected erythrocytes from beta thalassaemia heterozygotes is unknown, but clearly in these individuals there is scope for some of the effects of severe beta thalassaemia, such as oxidative damage from unpaired alpha globin chains, alterations in biophysical properties and hydration state^14^, and/or a reduction in the sialic acid content of glycophorins^15,16^ to be present, albeit to a more limited extent.

Impaired growth of malaria parasites in beta thalassaemic cells is one of the main candidate malaria protective mechanisms explored so far in *in vitro* studies. A second, not mutually exclusive potential protective mechanism is an enhanced removal of parasitised erythrocytes by the host immune system in beta thalassaemic individuals. Finally, it is possible that beta thalassaemia does not affect parasitaemia per se, but that it instead reduces the chance of severe malaria syndromes, such as cerebral malaria or severe malarial anaemia, developing.

Studies into these mechanisms have produced mixed results. Some authors have observed impaired *P. falciparum* growth in heterozygous beta thalassaemic erythrocytes^17^ while others have found no difference^18,19^. Beta thalassaemic erythrocytes are particularly sensitive to oxidant stress, and parasite growth seems to be particularly inhibited if erythrocytes are exposed to higher oxygen levels (25-30%)^20^. “Impaired growth” includes several sub-categories of possible mechanisms: blocking of invasion, deficiency in intracellular parasite development, or deficiencies in erythrocyte rupture and merozoite release. A study which examined parasite invasion separately from maturation found no impact of beta thalassaemia heterozygosity on either invasion or maturation^18^. In terms of the impact of beta thalassaemia heterozygosity on merozoite release, the only study to address this directly observed that on average, *P. falciparum* strain 3D7 produces four fewer merozoites per infected beta thalassaemic red blood cell compared to wild type red blood cells^21^.

The second potential protective mechanism, enhanced immune clearance, is supported by the observation of enhanced antibody binding to parasitised heterozygous beta thalassaemic cells *in vitro*^22^, and further evidenced by the observation that beta thalassaemia erythrocytes infected with ring-stage *P. falciparum* parasites are phagocytosed more than ring-stages infecting non-thalassaemic cells^19^. This phenomenon may be due to a double oxidative stress, and potentially increased aggregation of band 3^19^. Reduced lateral mobility of band 3 has been shown to increase rigidity of erythrocytes and confer resistance to malaria in the case of Southeast Asian Ovalocytosis^23,24^. The third potential protective mechanism, specific protection against severe malaria syndromes, has not been explored for beta thalassaemia.

Here, we conduct a detailed study of the erythrocyte membrane properties of beta thalassaemia heterozygotes and how these relate to invasion by different strains of *P. falciparum*. Erythrocyte membrane tension was recently shown to play a role in the malaria protective mechanism of the Dantu blood type^25^. We calculate membrane tension for beta thalassaemic erythrocytes, and investigate how this and other biophysical properties impact the wrapping energy required for merozoites to invade. We also investigate the membrane expression of proteins of known or potential importance to *P. falciparum* invasion. We observed significant changes in the membrane tension and bending properties of beta thalassaemic erythrocytes compared to wild type controls. However, whilst the change in tension acted to increase wrapping energy for merozoite invasion, the change in bending modulus acted to decrease wrapping energy for merozoite invasion, and overall we did not observe any *P. falciparum* invasion deficiency. We observed an increase in expression of basigin, an essential receptor for *P. falciparum* invasion, on beta thalassaemic cells compared to wild type controls, and we speculate that this change in expression level could also act against any *P. falciparum* invasion defect for beta thalassaemia.

## Results

### (i) Beta thalassaemia heterozygosity significantly affects erythrocyte membrane tension and bending modulus, but does not significantly affect the wrapping energy required for merozoite invasion of erythrocytes

The blood samples used in this study were sent from the King’s College Hospital to Cambridge for experiments in two different batches in 2018 and 2019, respectively (see Methods). We improved the method of transporting them to Cambridge in 2019, meaning that samples used in 2019 were fresher upon arrival than samples used in 2018. Hence, we present the results for samples from each year separately throughout the manuscript. Biophysical parameters of erythrocytes from beta thalassaemia heterozygotes, along with erythrocytes from controls collected on the same day and in the same clinic, were measured using cell contour flickering as described^26^.

For each cell, we measure two biophysical parameters: its bending modulus and its tension. Both measure the energy needed to deform the cell membrane from its equilibrium shape. The cell’s bending modulus is the energy required to bend an area of the cell membrane, whereas the cell’s tension measures the energy required to stretch its effective surface area (**Supplemental Methods S1.1**). Erythrocytes from beta thalassaemia heterozygotes tend to have a higher membrane tension (**Figure 1a**) and a lower bending modulus (**Figure 1b**) than their matched controls. The pattern is most consistent in the fresher 2019 samples. 2019 samples (n=3 pairs) had lower tension values in general than those from 2018, suggesting that time spent outside of the body increases the tension of red blood cells generally, as founded in literature^27^. Beta thalassaemic status is a significant predictor of tension (p = 0.0259) in a mixed model accounting for both individual patient and day of sample collection (i.e. conditions experienced by the cells) as random effects. The average increase in tension associated with beta thalassaemia heterozygosity is 4.78 × 10^−7^ N m^-1^ (95% CI 0.68, 8.84 × 10^−7^ N m^-1^). Likewise, beta thalassaemic status is a significant predictor of bending modulus (p = 0.0144) in a mixed model accounting for both individual patient and day of sample collection as random effects. The average change in bending modulus associated with beta thalassaemia heterozygosity is -4.08 × 10^−20^ J (95% CI -0.957, -7.11× 10^−20^ J).

**Figure 1:**
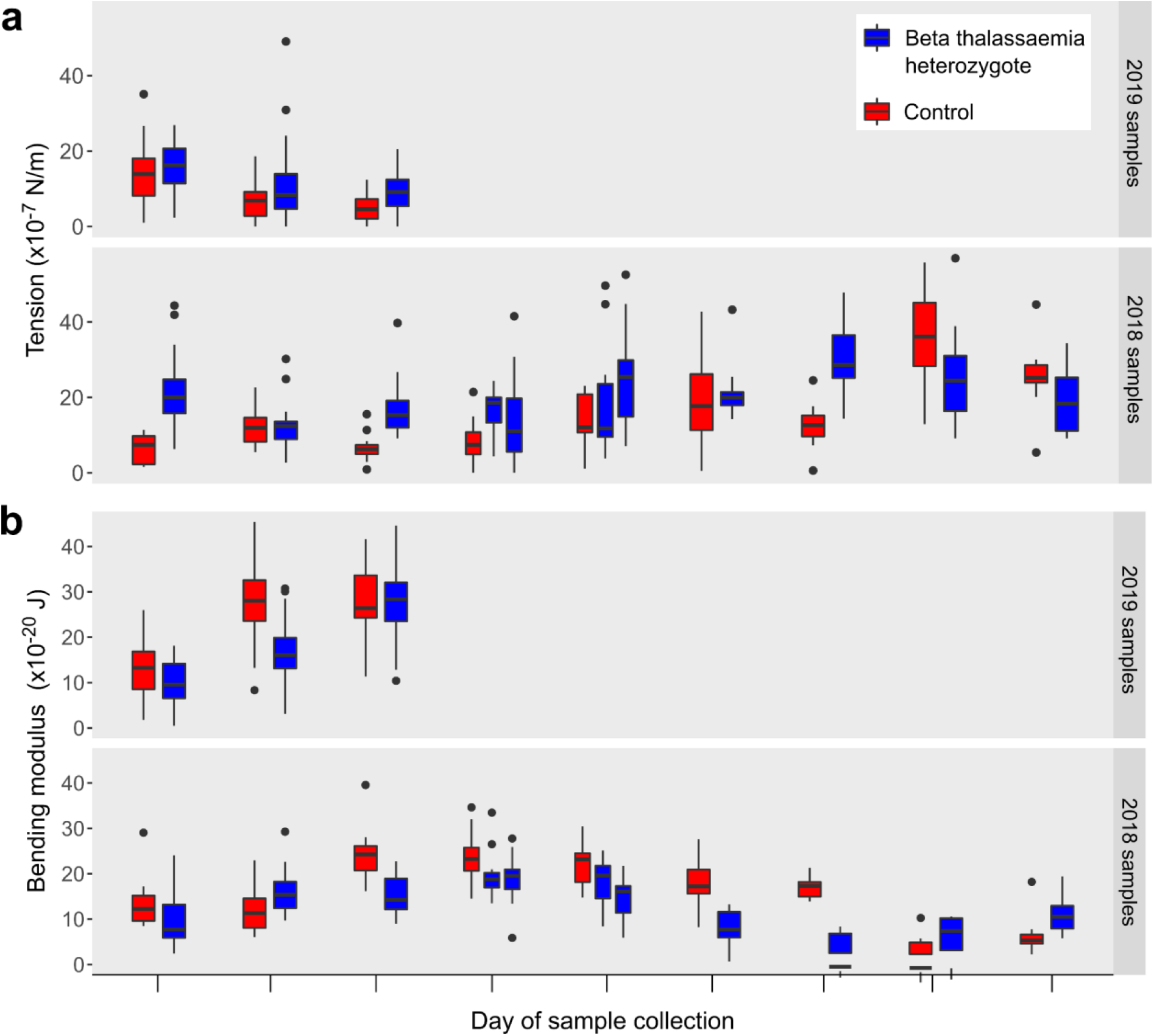
Membrane tension and bending modulus in beta thalassaemic erythrocytes. The boxplots in panels **a** and **b** display the erythrocyte membrane tension (**a**) or bending modulus (**b**) measured for 14 beta thalassaemia heterozygotes (blue boxplots) and 12 control non beta thalassaemic samples (red boxplots). Samples subject to the same conditions (i.e. gathered on the same day and tested as part of the same batch) are grouped together. In two cases, two beta thalassaemic samples were gathered on the same day and a single control non-beta thalassaemic sample from that day was used for both. The number of cells measured to produce the results in each boxplot range between 7 and 50, and 2019 samples spent less time in transit than 2018 samples and hence were fresher (see Methods).

We used the tension and bending modulus measurements, as well as previously calculated merozoite and erythrocyte dimensions^28^, to calculate for the first time an approximate energy of erythrocyte deformation, accounting for the stretch and curvature of the cell membrane as it “wraps” a merozoite (**Figure 2a**). Beta thalassaemic cells tend to have higher wrapping energies than their matched controls (**Figure 2b**), meaning it would theoretically take more energy for the merozoite to deform the membrane for a successful invasion event. The average difference is 100 x10^−20^ J according to a mixed model accounting for both individual patient and day of sample collection as random effects. However, the 95% confidence interval for this estimate ranges between -14 and 203 x10^−20^ J, and the likelihood ratio method we used to determine significance in our mixed models (see Methods), did not conclude that there was a significant difference in wrapping energy between beta thalassaemia heterozygotes and their matched controls at α=0.05 (p =0.05347). The significant differences we observed in both tension and bending modulus therefore do not translate into a significant difference in wrapping energy. This may in part be because our observed differences in tension and bending modulus act in opposite directions. As illustrated in **Figure 1**, beta thalassaemia heterozygous erythrocytes have higher tensions (increasing wrapping energy) but lower bending modulus values (decreasing wrapping energy) than their matched controls.

**Figure 2:**
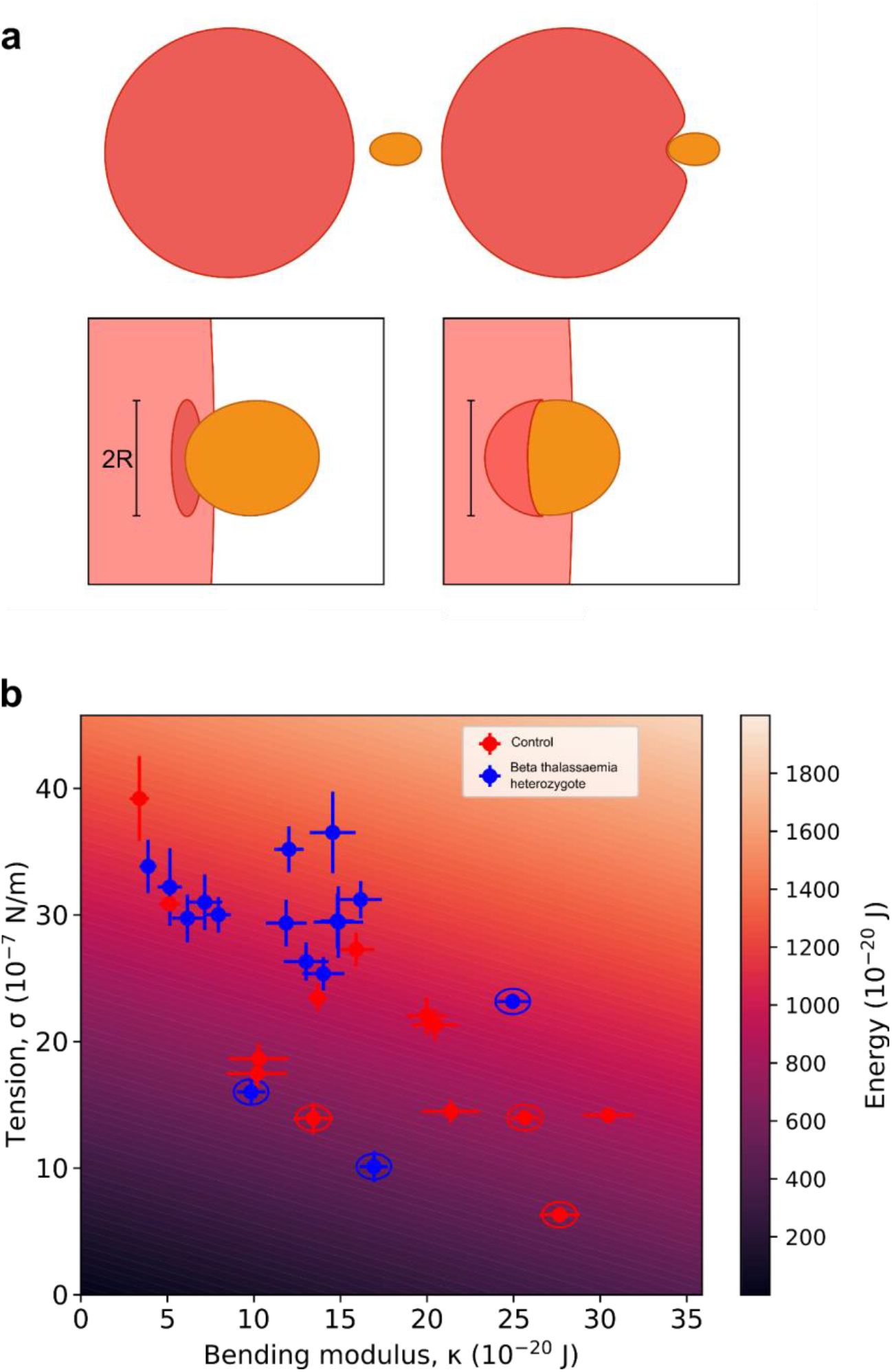
Wrapping energy in beta thalassaemic erythrocytes. Panel **a** shows a cartoon of the merozoite (orange) of radius R approaching the erythrocyte (red) and the subsequent deformation. In the top, sequentially from left to right, we see that the merozoite attachment precedes a localised wrapping of the erythrocyte membrane around it. In the lower two panels, the same process is magnified and the geometry emphasised. The patch of membrane which is involved in wrapping is modelled first as a disc of radius R, and following wrapping as a half-sphere of the same radius. Such a deformation requires work, as it involves both bending and stretching of the erythrocyte membrane. Panel **b** shows the wrapping energy, which is a function of both bending modulus and stiffness. The heatmap shows the form of this function, with the intensity of background shading representing different wrapping energy, which changes dynamically as tension and bending modulus change. The wrapping energy was also calculated for each individual sample using the data generated in **Figure 1**. The error bars are the standard error of the mean for each sample. 2019 samples are circled.

As a control for this novel approach we calculated wrapping energies for Dantu homozygous erythrocytes using the data in our previous paper Kariuki *et al*.^24^, and found that, unlike for beta thalassaemic cells, an increase in tension drives significantly higher wrapping energies for Dantu homozygous erythrocytes (**Supplementary Figure S1a**), correlating with our observation in that work that *P. falciparum* invasion into Dantu homozygote erythrocytes is much less efficient than into non-Dantu erythrocytes. Unlike beta thalassaemic cells, the bending modulus in Dantu blood group erythrocytes does not change with respect to non-Dantu^24^.

### (ii) Differences in tension or wrapping energy do not drive consistent differences in *P. falciparum* invasion between beta thalassaemic and non-beta thalassaemic erythrocytes

To establish whether these changes in biophysical properties were associated with any difference in the ability of *P. falciparum* to invade beta thalassaemic cells, we performed assays using four different *P. falciparum* strains selected for their different invasion pathway usage (3D7, 7G8, Dd2 and GB4; based on collection site and/or genome analysis these strains originate from West Africa (3D7, GB4), Brazil (7G8) and Southeast Asia (Dd2) respectively). We observed no consistent pattern of lower *P. falciparum* invasion for beta thalassaemia heterozygote erythrocytes measured by either an invasion assay in which each cell type was invaded separately (**Figure 3a**), or a preference assay in which beta thalassaemic cells and their matched control cells were mixed together and thus exposed to exactly the same invasion conditions (**Figure 3b**). For Dd2 the preference assay indicated that beta thalassaemic cells were significantly less efficiently invaded than control cells (**Figure 2b**), but there was no such pattern in the corresponding invasion assay, which reduces confidence in the significance of the difference. We also used live imaging video to observe 3D7 invasion events in a subset of samples. While there was an overall higher proportion of successful invasion events in control cells, we again did not observe consistent differences between beta thalassaemia heterozygote/control pairs (**Supplementary Figure S2**).

**Figure 3:**
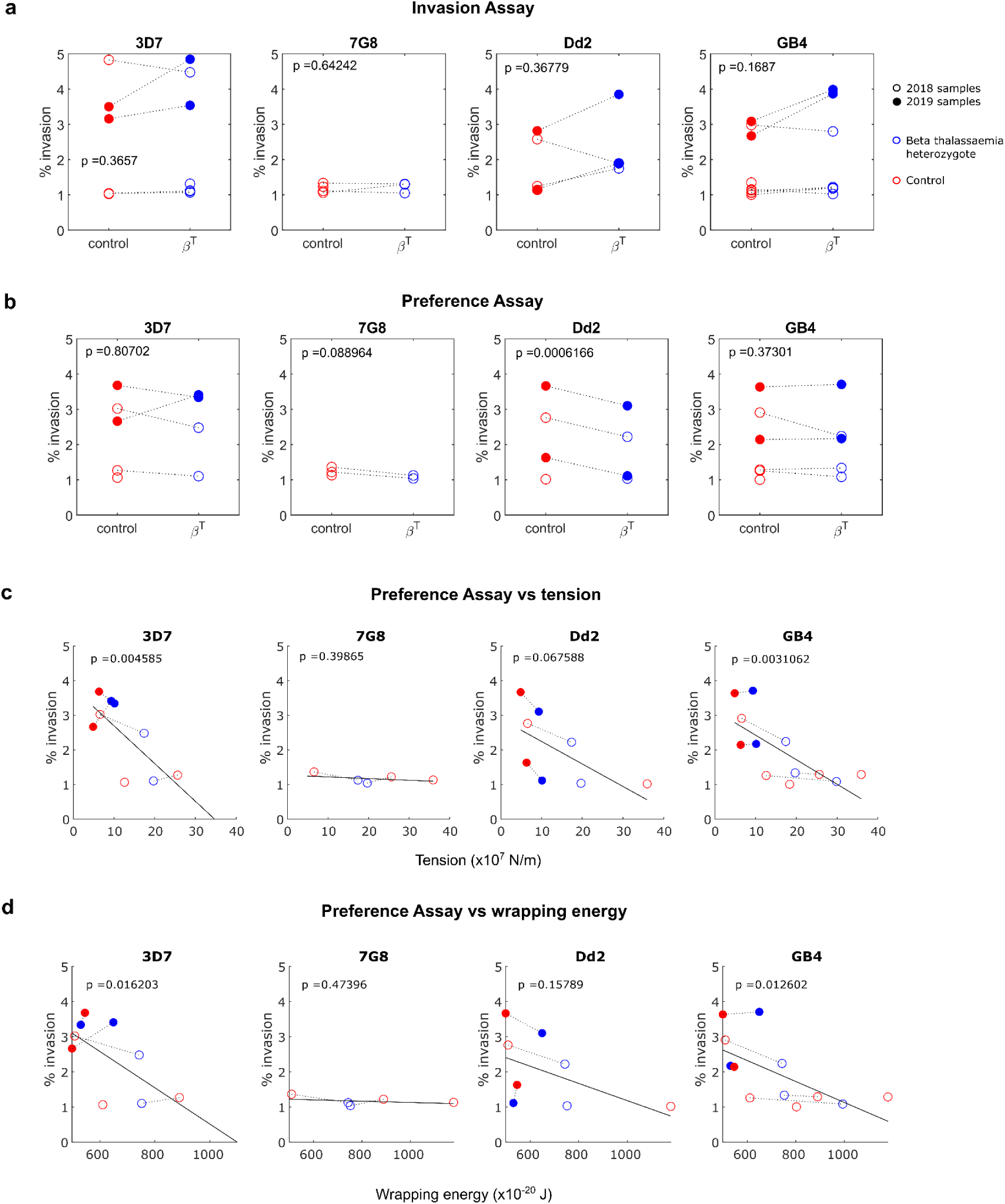
*P. falciparum* invasion and preference assays for beta thalassaemic and control erythrocytes. Panels **a** and **b** illustrate the results of two different forms of invasion assay (see Methods), one in which each cell type is invaded separately (panel **a**, “invasion assay”) and one in which both cell types are mixed together and subject to exactly the same invasion conditions (panel **b**, “preference assay”). Each marker is the mean of three technical replicates. Only results where the % invasion was greater than or equal to 1% have been shown, since % invasion results lower than this are likely to be unreliable. Dashed lines join beta thalassaemia heterozygous samples with their matched controls; if a sample does not have a pair this is because the matched sample had a % invasion <1%. 2018 samples are illustrated with open markers and 2019 samples with filled markers. The p values in each sub plot of panels **a** and **b** are the results of paired T tests testing the null hypothesis of no difference in either % invasion or preference score between the matched pairs. Panel **c** plots mean tension values for each sample (from the data in **Figure 1b**) against preference assay results, and panel **d** plots mean wrapping energy per sample against preference assay results. As in panels **a** and **b**, dashed lines connect beta thalassaemic samples with their matched controls. The solid lines in panels **c** and **d** are least squares regression lines for the relationship between either mean tension per sample (panel **c**) or mean wrapping energy per sample (panel **d**) and % invasion (disregarding beta thalassaemic status). The p values in panels **c** and **d** test whether the gradients of these least squares regression lines are significantly different from zero.

Comparing the average membrane tension of each sample with the invasion data revealed a negative relationship (i.e. higher tensions were associated with lower invasion), which was statistically significant for the strains for which we had the most datapoints (3D7 and GB4), supporting our previous findings of a link between tension and invasion (**Figure 3c**, solid trend lines). However, within each beta thalassaemic/matched control pair (**Figure 3c**, dashed lines) an increase in tension did not consistently predict a decrease in % invasion.

Increases in wrapping energy, a parameter estimated for the first time in this work, were associated with reduced invasion (**Figure 3d**, solid trend lines), with significant associations for the 3D7 and GB4 strains. However, within each beta thalassaemic/control pair there were two cases where a beta thalassaemic sample had higher wrapping energy than its matched control, but had similar or even higher invasion, emphasising that there are outliers within the overall trends (**Figure 3d**, dashed lines). Overall, it appears that the differences in tension or wrapping energy between beta thalassaemia heterozygotes and control cells may not be large enough to consistently drive differences in % invasion.

In our Dantu study^24^ we measured invasion events and biophysical properties of individual cells simultaneously, which was not performed here. However, applying our wrapping energy calculations to that previously collected Dantu dataset, showed that cells which were invaded had significantly lower wrapping energies than those which failed to be invaded (**Supplementary Figure S1b**). The results in **Figure 3** are limited by the fact that so many of our invasion assay results produced % invasions lower than 1% and had to be discarded, so our beta thalassaemia invasion dataset is very small.

### (iii) *P. falciparum* merozoite adhesion force may be slightly lower in beta thalassaemic erythrocytes

We measured the strength of the interaction between 3D7 or Dd2 *P. falciparum* merozoites and beta thalassaemia heterozygotes or matched control non-beta thalassaemic samples using the optical tweezer extension assay (see Methods and ^29^). Note that to calculate the adhesion force, the stiffness of the erythrocyte membrane is taken into consideration, and at near-zero strain rate (<0.1 s^-1^) it was previously measured^28,30^ at 20 pN *μ*m^-1^. We verified that stiffness values are comparable in beta thalassaemic and control cells (**Supplementary Figure S3**), and therefore applied the 20 pN *μ*m^-1^ stiffness value to all cells in this analysis. As shown in **Figure 4**, the median adhesion force values are lower in the beta thalassaemic sample in 5/6 of the pairs, but the difference was not significant, with a p value of 0.2188 in a simple sign test of the null hypothesis that the adhesion force values are equally likely to be higher or lower in the beta thalassaemic cells compared to the control cells. Unfortunately, we did not obtain enough results from different individuals to fit a reliable mixed model accounting for both individual patient and day of sample collection as random effects.

**Figure 4:**
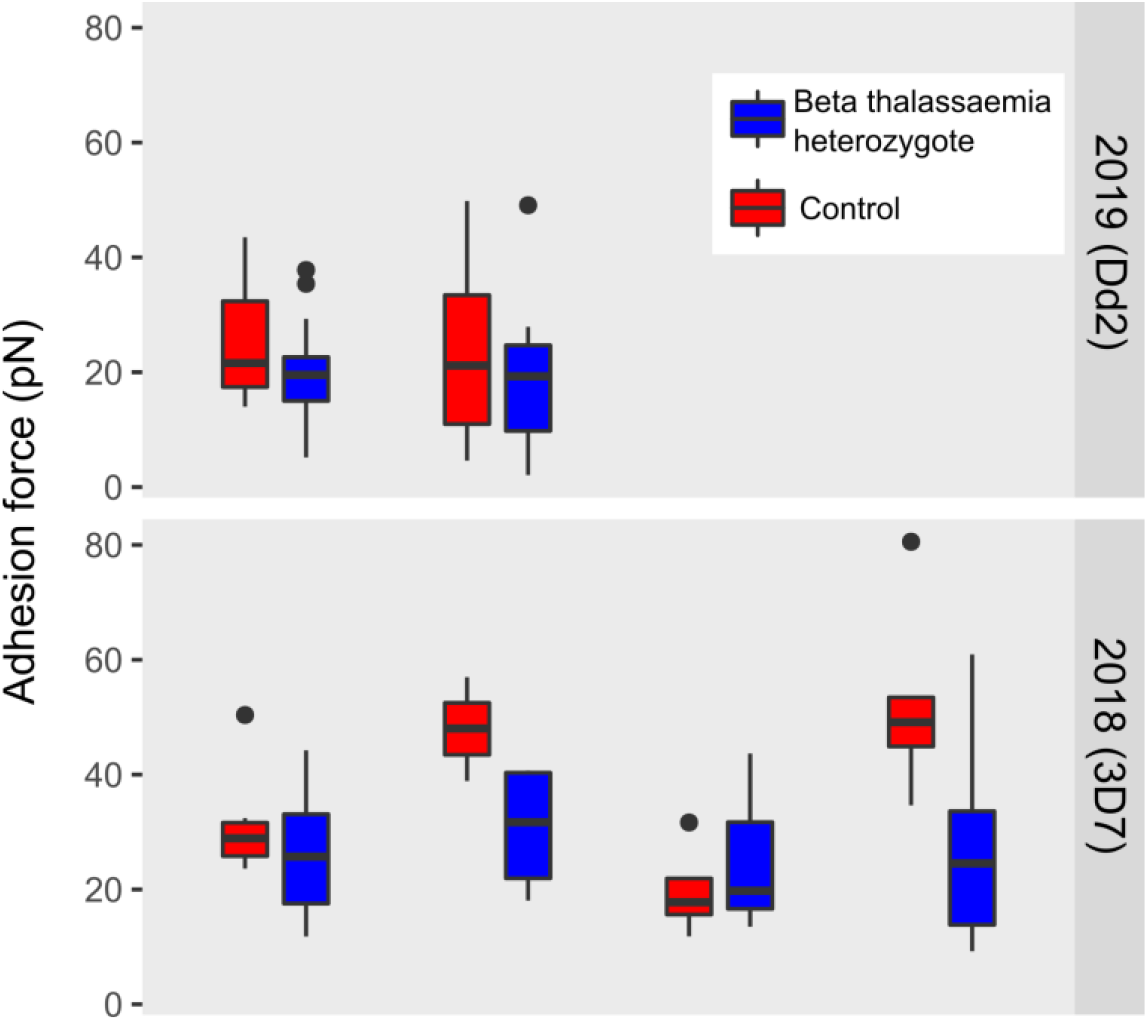
*P. falciparum* adhesion force for beta thalassaemic erythrocytes. The boxplots display merozoite adhesion force measured for 6 beta thalassaemic heterozygotes (blue boxplots) and 6 control non beta thalassaemic samples (red boxplots). Samples subject to the same conditions (i.e. gathered on the same day) are grouped together. The number of cells measured to produce the results in each boxplot range between 3 and 15. Parasite strains used were 3D7 in 2018 and Dd2 in 2019.

### (iv) Characterisation of erythrocyte surface proteins in the membranes of beta thalassaemic cells

We used flow cytometry to quantitate the erythrocyte membrane expression levels of 11 proteins with a known or possible role in *P. falciparum* invasion: complement receptor 1; CD44 antigen; integrin alpha 4; complement decay accelerating factor; transferrin receptor protein; basigin; band 3 anion transport protein; Duffy antigen/chemokine receptor; glycophorin A; glycophorin B, and glycophorin C. Beta thalassaemic cells appeared to have higher expression of basigin, and lower expression of band 3, Duffy antigen and glycophorin A than non-beta thalassaemic controls (**Figure 5a**); however if we apply a stringent Bonferroni correction for multiple comparisons (where in this case, significance at α=0.05 requires p≤0.0045), none of these proteins display a significant difference in expression level.

**Figure 5:**
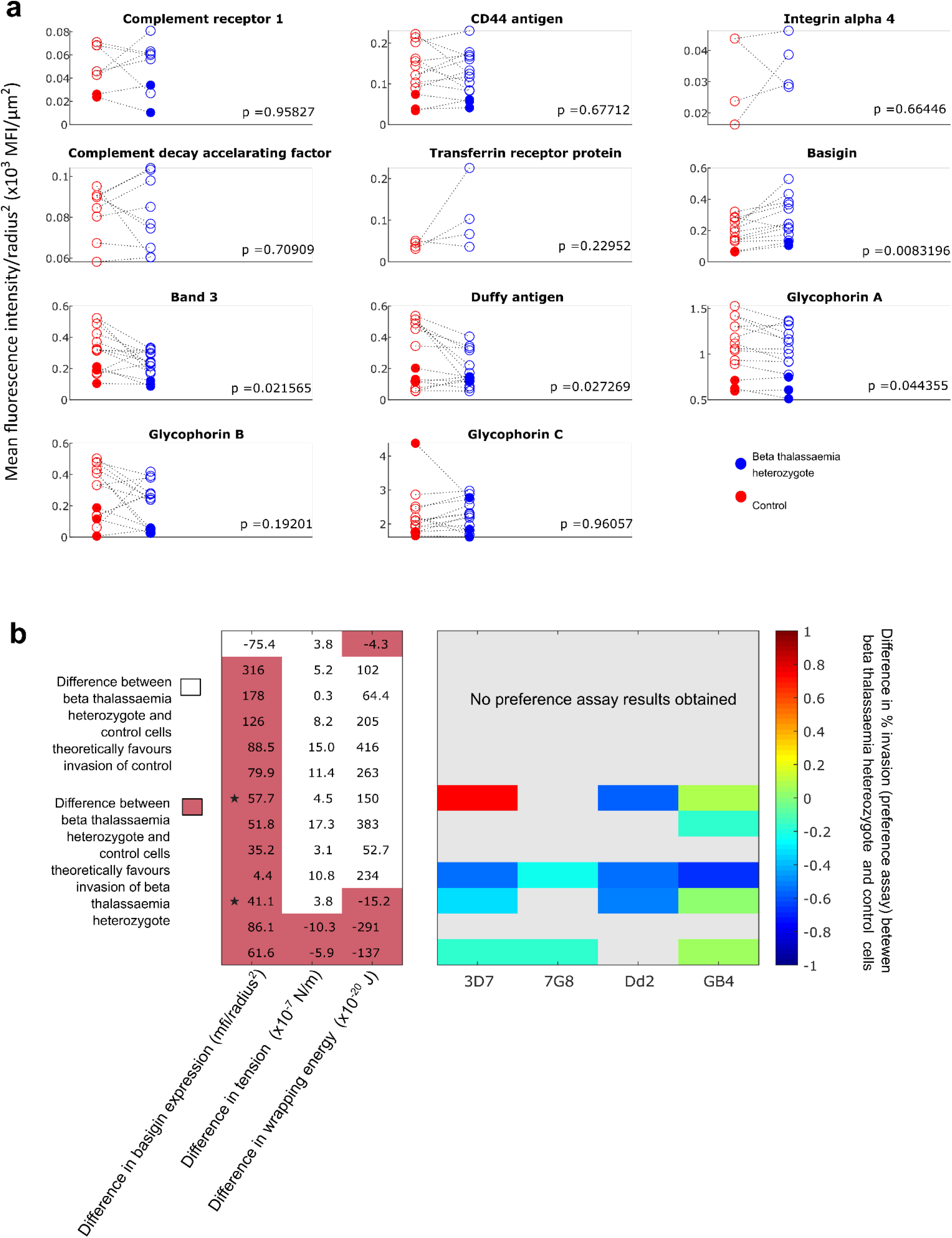
Expression of potential *P. falciparum* invasion proteins in the membrane of beta thalassaemic and non-beta thalassaemic erythrocytes, and possible impact on invasion. Panel **a** displays the expression levels of 11 different potential *P. falciparum* invasion proteins, in beta thalassaemia heterozygous red blood cells and matched control red blood cells. Expression levels are calculated as the mean fluorescence intensity obtained for each sample (corrected for background noise - see Methods), divided by the square of the mean erythrocyte radius calculated for each sample. Dividing fluorescence intensity by the square of the erythrocyte radius corrects for the fact that beta thalassaemia heterozygotes tend to have smaller red blood cells than non-beta thalassaemic individuals. Each marker is the mean of three technical replicates. Dashed lines connect beta thalassaemia heterozygote samples to their matched controls. 2018 samples are shown using empty markers, 2019 samples using filled markers. The p values shown in each subplot are the results of paired T tests testing the null hypothesis of no difference in expression level between the matched pairs. Panel **b** summarises the differences in mean basigin expression level, mean membrane tension, and mean wrapping energy between beta thalassaemic cells and control cells (specifically: beta thalassaemic value – control value), for 13 matched pairs where all 3 pieces of information are available (each row = one pair). Differences which theoretically favour the invasion of beta thalassaemia heterozygous cells rather than control cells are indicated with a white background and differences which theoretically favour the invasion of control cells rather than beta thalassaemia heterozygous cells are indicated with a red background. We also summarise the difference in preference assay results for each beta thalassaemic/control pair, for each of the strains of parasite tested (the same data as in **Figure 2b**, expressed as the beta thalassaemic preference score minus the control preference score). The rows in panel **b** which correspond to 2019 samples have been indicated with a star.

Basigin is known to be an essential invasion receptor for all *P. falciparum* strains studied^31^. Since so many of our beta thalassaemic samples had higher basigin expression than their matched controls, we analysed whether a combination of (i) basigin expression, (ii) membrane tension, and/or (iii) wrapping energy could account for the invasion properties of our samples. As shown in **Figure 5b**, *none* of the pairs of samples consistently displayed traits which would theoretically favour invasion of control cells rather than beta thalassaemic cells. There were no pairs of samples with a lower level of basigin expression in the beta thalassaemic cells than in the control cells, alongside a higher wrapping energy for the beta thalassaemic cells than the control cells, and a higher predicted proportion of invadable control cells. We cannot be certain if the differences in basigin expression observed here are functionally important, but the direction of the patterns observed opens up the possibility that the impact of wrapping energy or tension on the invasion of beta thalassaemic cells is being counteracted by a change in basigin expression. Such a phenomenon could account for the lack of consistent effects of beta thalassaemic status on *P. falciparum* invasion.

## Discussion

We have conducted a detailed study of the properties of heterozygous beta thalassaemic erythrocytes with respect to *P. falciparum* invasion, including the first calculations of merozoite wrapping energy as a potential explanatory factor in the invasion process. We found a range of invasion-relevant changes in heterozygous beta thalassaemic cells (higher tensions and bending modulus; changes in basigin expression) but observed no significant changes in the percentage of invaded beta thalassaemic cells in invasion or preference assays. Our results thus suggest there is no invasion defect in heterozygous beta thalassaemia, which concurs with the conclusions of a previous study by Ayi *et al*.^18^. However, our results raise the possibility that the lack of an invasion defect in heterozygous beta thalassaemia arises from a complex combination of altered membrane properties, including some changes which act to decrease the probability of successful *P. falciparum* invasion (higher tension) and others which act to increase it (lower bending modulus; higher basigin expression).

We have previously shown that the Dantu blood group, a glycophorin mutation which reaches allele frequencies of up to 10% in Kilifi, Kenya, is associated with increased erythrocyte membrane tension, and that this increase in tension leads to fewer successful merozoite invasion events^24,32^. The average increase in erythrocyte tension we observe for beta thalassaemic heterozygotes (4.78 × 10^−7^ N m^-1^) is comparable to that observed between non-Dantu individuals and Dantu homozygotes (2.8 × 10^−7^ N m^-1^), but does not seem to decrease invasion. This apparent contradiction is resolved when we consider bending modulus and wrapping energy. Beta thalassaemic cells display a higher bending modulus than wild-type cells whereas Dantu homozygotes have no such change. The wrapping energy increase associated with beta thalassaemia heterozygosity is 100 x10^−20^ J (which we found was not a significant difference in our analysis), but the wrapping energy increase associated with Dantu homozygosity is 258 × 10^−20^ J (**Supplementary figure S1a**). This greater wrapping energy increase could account for an invasion deficit being seen in Dantu erythrocytes but not in beta thalassaemia heterozygous erythrocytes.

A further factor complicating the invasion of beta thalassaemia heterozygous erythrocytes is the expression of *P. falciparum* invasion receptors in the erythrocyte cell membrane. We observed increased basigin expression in beta thalassaemia heterozygous cells as opposed to matched non-beta-thalassaemic controls. Basigin is an essential receptor for *P. falciparum* invasion. Interfering with the interaction between basigin and its parasite ligand *Pf*Rh5 by targeting either the host or parasite protein results in substantial invasion defects^30^. However, the impact of basigin expression levels on invasion in general, and the potential impact of changes such as those we report in **Figure 5**, is unknown. Is there a correlation between basigin expression and invasion success, or is there simply a threshold level of basigin required for invasion? If there *is* a correlation between the levels of basigin present on an erythrocyte and invasion success, increased basigin expression on beta thalassaemic cells provides an additional reason that beta thalassaemic cells should display no invasion defect, despite their higher tension relative to control red blood cells. The red blood cell population of a person with thalassaemia is skewed towards younger red blood cells, and it has been found that immature red blood cells, reticulocytes, express more basigin than older cells^33,34^, but more work is needed to establish what may drive higher basigin levels in beta thalassaemia.

If erythrocytes from beta thalassaemia heterozygotes are invaded at a normal rate by *P. falciparum*, what is the advantage of beta thalassaemia heterozygosity against malaria? As noted in the introduction, one possibility which has so far received little attention is that beta thalassaemia may protect against the development of severe malaria syndromes. Alpha thalassaemia, whilst a different condition, potentially provides an informative comparison. *P. falciparum* infected alpha thalassaemic red blood cells display reduced cytoadherence to endothelial tissues^35,36^, a phenomenon which is likely to reduce severe malaria pathology. The exact mechanism behind this reduced cytoadherence is not fully understood. Some studies find reduced expression of *Pf*emp1, the major *P. falciparum* cytoadherence ligand, on the surface of infected alpha thalassaemic erythrocytes^34^ but others do not^35^. Understanding how/whether the beta thalassaemia-associated erythrocyte membrane alterations we have demonstrated here might impact cytoadherence in *P. falciparum* infected beta thalassaemic cells would be highly informative.Our study had some limitations, in particular the freshness of the 2018 blood samples and the low % invasions observed in some samples which limited our invasion assay results. However, since we observed our phenomena of interest (increased tension, decreased bending modulus, increased basigin expression) in the fresher 2019 samples as well as the less fresh 2018 samples, and since each beta thalassaemic sample had a matched control which experienced the same conditions, allowing us to statistically account for effects of day of sampling, we remain confident in our observations overall.

The *P. falciparum* invasion properties of beta thalassaemic cells, where changes to the erythrocyte membrane may counteract each other to result in no overall invasion defect, add to an overall picture in which widespread human adaptations to *P. falciparum* tend not to convincingly reduce parasite invasion in *in vitro* studies. Glucose-6-phosphate dehydrogenase (G6PD) deficiency and alpha thalassaemia are found in all old-world malarious regions, but neither G6PD deficiency^37^ nor alpha thalassaemia^38^ reduce invasion. Sickle cell trait may have a small impact on invasion but this is not observed in all studies^39,18^. Adaptations which reduce invasion are more limited in their distribution: Southeast Asian Ovalocytosis may reduce invasion of strains which use specific invasion pathways^40^, but is limited to Malaysia and Papua New Guinea, and the Dantu blood group which, as noted above, reduces parasite invasion, is limited to a relatively small region of East Africa.

A recent theoretical evolutionary study may shed some light on these pattern^41^. If it is possible to gain adaptive immunity against the worst impacts of *P. falciparum* malaria after experiencing relatively few blood stage infections, this limits the advantage of a mutation which partially blocks *P. falciparum* invasion. Indeed, in a competition between two adaptations, both of which reduce the probability of severe disease occurring but only one of which simultaneously reduces the probability of invasion, this model predicts that the adaptation which *does* protect against severe disease but *does not* reduce invasion will spread more quickly. It may be that the success of beta thalassaemia, and other globally widespread *P. falciparum* adaptations is because of, rather than in spite of, their limited impact on *P. falciparum* invasion.

## Materials and Methods

### Sample collection

Anonymised fresh blood samples were obtained after they were discarded from the haemoglobinopathy diagnostics service of King’s College Hospital London (KCH), with ethical approval (approved by the Camden and King’s Cross REC, reference 13/LO/0728). The beta thalassaemic status (or control status) of these samples had been diagnosed by haemoglobin high performance liquid chromatography (HPLC). Heterozygous beta thalassaemia was diagnosed if HbA_2_ levels exceeded 4%. No genetic analysis was done, but the presence of HbS, HbC or HbE was ruled out for all samples as part of the screening. Co-inherited alpha thalassaemia or G6PD deficiency was still possible in any of the samples. 16 samples from beta thalassaemia heterozygous individuals and 13 control samples (matched by day of sample acquisition; on some days more than one beta thalassaemic sample was available, and just one matching control sample was provided) were shipped to the Wellcome Sanger Institute and to the Cavendish Laboratory, where all the experiments were part of a blind study. In 2018 (13 beta thalassaemic samples, 10 controls), we allowed the use of any sample discarded within the previous 7 days; samples were refrigerated whilst stored at KCH and sent to Cambridge using Royal Mail next day delivery, thus no sample should have been more than 8 days old when first used. One set of samples was delayed in transit and therefore not included in any analysis here. In 2019, samples were dispatched from KCH to Cambridge in a chilled package via a courier service, within 24 hours of having been drawn from the patient, thus samples arrived in Cambridge for experiments to begin within 48 hours of having been drawn from the patient.

### *In vitro* culture of *P. falciparum* parasites

All *P. falciparum* parasite strains used in this study (3D7, Dd2, 7G8, GB4) were routinely cultured in human O-erythrocytes (NHS Blood and Transplant, Cambridge, UK) at 3% haematocrit (Hct) in RPMI 1640 medium with 25 mM Hepes, 20 mM glucose, and 25 μg mL^-1^ gentamicin containing 10% Albumax at 37°C (complete medium), under an atmosphere of 1% O_2_, 3% CO_2_, and 96% N_2_ (BOC, Guildford, UK). Parasite cultures were synchronised on early ring stages with 5% D-sorbitol (Sigma-Aldrich, Dorset, UK). Use of erythrocytes from human donors for *P. falciparum* culture was approved by the NHS Cambridgeshire 4 Research Ethics Committee.

### Invasion and preference assays

Erythrocytes were stained with two concentrations of CellTrace Far Red Cell Proliferation kit (Invitrogen, UK) – 4 µM Heterozygotes β-thalassaemic cells (β-thal) and 16 µM in non-β-thal cells - as described here^24^. After 2 hour incubation at 37°C under rotation, the stained cells were washed and resuspended to 2% Hct with complete medium. Parasite cultures containing mostly schizont forms at 2-3% parasitaemia were pooled with equal volumes of erythrocytes from each genotype group in a 100 μl final volume. Thus, in the preference assay, 33 µl schizont culture + 33 µl β-thal + 33µl non-β-thal erythrocytes were pulled together in the same well in round bottom 96-well plates. While in the invasion assay, to evaluate whether the different concentrations of the dye could affect parasite growth, schizonts were mixed with stained erythrocytes from each genotype group in individual wells in a 1:1 ratio (50 µl schizont culture + 50 µl stained RBCs). The samples were incubated for 24 hours, then treated with 0.5 mg mL^-1^ ribonuclease A (Sigma Aldrich, UK) in PBS for 1 hour at 37°C to remove any trace of RNA^42^. Infected cells were stained with 2x SYBR Green I DNA dye (Invitrogen, Paisley, UK) in PBS for 1 hour at 37°C. Stained samples were examined with a Cytoflex S flow cytometer (Beckman Coulter, UK). SYBR Green I was excited by a blue laser and detected by a 530/30 filter. CellTrace Far Red was excited by a red laser and detected by a 660/20 filter. 50,000 events were collected for each sample. The data collected were then further analysed with FlowJo (Tree Star, Ashland, Oregon) to obtain the percentage of parasitised erythrocytes within each genotype group. All experiments were carried out in triplicate.

### Characterization of erythrocyte membrane by flow cytometry

A panel of antibodies was selected against 11 antigens that have been confirmed to be or could be potentially involved in cell adhesion and parasite invasion. Each blood sample was diluted at 0.5% Hct, washed twice with PBS and incubated in primary mouse monoclonal antibodies for 1 hour at 37°C. The antibodies used were: anti-CD35-APC (CR1, Thermofisher, 1:50); antiCD44-BRIC 222-FITC (1:100, IBGRL); anti-CD55-BRIC-216-FITC (1:500, IBGRL); Transferrin R: anti-CD71-PE (1:100, ThermoFisher); Basigin: anti-CD147-FITC (1:100, ThermoFisher); Band3: anti-CD233-BRIC6-FITC (1:1000, IBGRL); Duffy antigen: anti-CD234-APC (1:100, Milteny Biotec); GYPA: CD235a-BRIC 256-FITC (1:1000, IBGRL); GYPC: anti-CD236R-BRIC10-FITC (1:1000, IBGRL). For detection of GYPB, first cells were incubated with an anti-GYPB (1:100, rabbit polyclonal antibody, Abcam), then washed twice with PBS and then incubated with a goat-anti-rabbit AlexFluor488 labelled antibody. After incubation, cells were washed twice in PBS and analysed on a Cytoflex S flow cytometer. A control was carried out on cells with no antibody added to measure background fluorescence. Data were analysed using FlowJo Software (Treestar, Ashland, Oregon). All experiments were carried out in triplicate. Before statistical analysis, the average of the 3 control “no antibody” fluorescence readings were subtracted from the average of the 3 fluorescence readings for each sample, to generate a noise-corrected value. Any cases where the resulting noise-corrected value was less than maximum value of the original 3 “no antibody” readings were discarded, on the basis that this measurement was insufficiently different from the background noise.

### Live imaging of invasion and measurement of merozoite detachment force

Highly concentrated (97%) parasites (strains 3D7 and Dd2) were isolated by magnetic separation at the schizont stage (LD columns, Miltenyi Biotec, UK) and re-suspended in complete medium with either beta thalassaemic or non-thalassaemic erythrocytes at 0.05% Hct. The resulting sample was loaded in a SecureSeal Hybridization Chamber (Sigma-Aldrich) attached to a glass cover slip coated with 10 µl solution of poly(l-lysine)-graft-poly(ethylene glycol) (PLL-g-PEG) (SuSoS AG, Dübendorf, Switzerland) at 0.5 mg/mL concentration and incubated for 30 minutes to prevent excessive adherence of cell proteins onto the coverslip. Live imaging of parasite egress and invasion was performed as described here^24^ using a Nikon Eclipse Ti-E inverted microscope (Nikon, Japan) with a 60X Plan Apo VC 1.20 NA water objective, kept at physiological temperature through a heated collar. A custom-built temperature control system was used to maintain the optimal culture temperature of 37°C throughout the experiments. Samples were placed in contact with a transparent glass heater driven by a PID temperature controller in a feedback loop with the thermocouple attached to the glass slide. Motorised functions of the microscope were controlled via custom software written in-house and focus was maintained throughout the experiments using the Nikon Perfect Focus system. Images were acquired in bright-field with red filter using a CMOS camera (model GS3-U3-23S6M-C, Point Grey Research/FLIR Integrated Imaging Solutions (Machine Vision), Ri Inc., Canada) at 60 frames/s, with pixel size corresponding to 0.0973 µm. The optical tweezer setup fits onto the Nikon microscope and consists of a solid-state pumped Nd:YAG laser (IRCL-2W-1064; CrystaLaser, Reno, NV) having 2 W optical output at a wavelength of 1064 nm. The laser beam was steered via a pair of acousto-optical deflectors (AA Opto-Electronic, Orsay, France) controlled by custom-built electronics that allow multiple trapping with subnanometer position resolution.

Merozoite invasion efficiency in live videos was calculated as the fraction of merozoites that contacted and successfully invaded erythrocytes divided by the number of all merozoites that contacted nearby cells post-egress. This definition takes into account the fact that in a chamber we can have several invasions when more merozoites invade the same erythrocyte.

Adhesive forces at the merozoite-erythrocyte interface were quantified by evaluating the elastic morphological response of the erythrocyte as it resisted merozoite detachment. An optical trap was used to move a merozoite just after it had egressed closed to an uninfected cell, anchored to the coverslip. Trapping durations were kept short (< 10 s) to minimise any detrimental effect of local heating due to the laser beam. Attachment between merozoite and erythrocyte was then allowed to take place. Then a second erythrocyte was delivered close to the merozoite to form a cell-merozoite-cell system. The maximal elongation of the unanchored erythrocyte before detachment was measured by pulling it away from the point of attachment with the merozoite. Finally, since erythrocytes behave mechanically as a linear spring in this regime^29^, there is a linear relationship between force applied and stretch of the erythrocyte along a single axis. Erythrocyte stiffness (*k*) is 20 pN μm^-1^ with c. 20% uncertainty due to intrinsic variation between erythrocytes^28^. The detachment force is *F = k ΔL*, where the elongation of the erythrocyte (*ΔL*) is the difference between erythrocyte length measured in the frame before detachment occurs (*L*_*max*_) and its length at rest (*L*_*0*_).

### Measurement of erythrocyte stiffness using optical tweezers

Following the protocol previously optimised in our group^29^, 5 μm carboxylated silica beads (Bangs Labs) were washed in Mes buffer (Sigma M1317) and functionalised with Lectin (Sigma M1317) and EDC (Sigma E1769) (4 mg mL^-1^) for 8 hours at 37°C, allowing them to stick to erythrocytes. The aforementioned optical tweezer setup was used to bring two beads to diametrically opposed sides of the erythrocyte. One of the traps is kept at a fixed position, while the other is moved by a distance *ΔLaser*. The displacement of the bead in the fixed trap, *Δx*, is obtained using a custom image analysis code written in Python, based on a correlation kernel. The elongation of the cell is given by *ΔL = ΔLaser - 2Δx*. The strain is given by *γ = ΔL/L*_*0*_. In our experiments, we kept the strain rate at 1 s^-1^.

### Erythrocyte membrane contour detection and flickering spectrometry

Beta-thalassaemic and non-beta thalassaemic erythrocytes were diluted into culture medium at 0.01% Hct, and their membrane thermal fluctuations were recorded with the set-up described above at high frame rate (fps < 500 frames per s) and short exposure time (0.8 ms) for 20 s, with care taken to record cells at their equator. The contour for each frame was tracked by an in-house Python program; the algorithm underpinning it was based on one previously described^25^. Another in-house Python program decomposes the radial component of each contour into its Fourier modes, and computes the average power spectrum for each cell. The measured average power-spectrum is well-described by^43^:

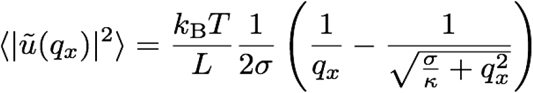

*u* is the amplitude of fluctuations, *q*_*x*_ is the mode, brackets denote averaging across all contours for a cell, *k*_*B*_ is the Boltzmann constant, *T* is the temperature, and *L* is the mean circumference of the erythrocyte contour. *σ* and *κ* are the tension and bending modulus, respectively, for which the same program fits the average power spectrum for mode numbers between 8 and 20.

### Erythrocyte membrane wrapping energy

The wrapping energy was calculated by considering changes to the free energy of the erythrocyte membrane due to its tension and the bending modulus. The sum of the two contributions are given by

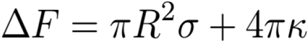

where *R* = 1 um is the radius of the merozoite^27^, *σ* is the tension, and *κ* is the bending modulus. The above equation is derived and developed in the **Supplementary Methods**.

### Statistical analysis

The biophysical measurements (tension, wrapping energy, adhesion force) involved taking repeated measures of erythrocytes from individual samples. Variation in these measurements is likely to derive from individual human differences as well as the way that samples were treated on a given day, hence we analysed these data using mixed models. All mixed models used individual sample identity and day of collection as random effects, and beta thalassaemia status of sample as a fixed effect. Mixed models were fitted in R version 4.0.3 [(R Core Team (2020). R: A language and environment for statistical computing. R Foundation for Statistical Computing, Vienna, Austria. URL https://www.R-project.org/.], using the lme4 package [https://cran.r-project.org/web/packages/lme4/citation.html], which also provided us with confidence intervals for effect sizes via the confint() function. Effect sizes and standard errors for the components of the mixed models were obtained using the arm package [https://cran.r-project.org/web/packages/arm/index.html]. We obtained a p value for the effect of beta thalassaemia status by fitting versions of each model with and without beta thalassaemia status as an explanatory factor, and conducting a likelihood ratio test to determine the difference between the models. P values quoted in the text for the mixed models are the p values from such likelihood ratio tests.

The invasion and preference assay results, and the antibody-based tests for membrane protein levels each involved carrying out three technical replicates of the relevant assay per sample. Each beta thalassaemic sample had a matched control sample, taken on the same day and subject to the same conditions. To analyse these data, we first averaged the technical replicates for each sample, and then used paired T tests to ascertain if there were systematic differences between beta thalassaemic samples and their matched controls.

Boxplots were plotted in R version 4.0.3 using ggplot2 [https://cran.r-project.org/web/packages/ggplot2/citation.html]; other plots were produced in Matlab version R2018b.

## Data Availability

The authors declare that the data supporting the findings of this study are available within this Manuscript, Supplementary Information and Methods.

## Acknowledgements

We thank Thomas Williams for providing insightful comments on the manuscript. V.I. was funded by the EPSRC, the Sackler fellowship, and the Wellcome Trust Junior Interdisciplinary Fellowship (Wellcome 20485/Z/16/Z). J.C.R. and his lab was funded by the Wellcome Trust (206194/Z/17/Z and 220266/Z/20/Z).

## Author contributions

V.I., A.M.-M., P.C., J.C.R., and B.S.P conceived and planned the experiments. D.C.R and J.N.B provided samples. V.I. and Y.-C.L. performed live invasion experiments, V.I. and G.N. performed and analysed flickering and optical tweezers experiments, A.M.-M. carried out erythrocyte preference invasion assay and erythrocyte membrane protein characterization by flow cytometry. G.N. carried out the analysis of the wrapping energy and B.S.P. analysed and plotted data. V.I. A.M.-M., G.N., P.C., J.C.R., and B.S.P. wrote the manuscript, and all authors reviewed and revised the final manuscript.

