## Supplementary Methods for "The erythrocyte membrane properties of beta thalassaemia heterozygotes and their consequences for *Plasmodium falciparum* invasion"

### S1 Wrapping energy of the membrane

In this section, we explain the form of the wrapping energy of the membrane by comparing the free energy of an erythrocyte under equilibrium with one that has a localised deformation.

The free-energy $F$ of an erythrocyte depends on the shape of the membrane. For our particular geometry, we can parametrize the shape of the membrane by $h$, its displacement relative to some equilibrium shape, the Monge parametrization. The free-energy can then be expressed by the Helfrich Hamiltonian [1, 2, 3].

$$F = \int_S \mathrm{d}S \left[ \frac{1}{2}\sigma(\nabla h)^2 + \frac{1}{2}\kappa(\nabla^2 h)^2 \right] \tag{S1}$$

$\kappa$ is the bending modulus, while $\sigma$ is the membrane's tension (See S1.1). The free energy is an integral over the surface $S$.

For small displacements this equation can be expressed as [3]:

$$F = \sigma A + 2\kappa \int_S \mathrm{d}S H^2 \tag{S2}$$

where $H$ is the mean curvature of the membrane.

We consider the deformation to be localized to a disc of radius $R$, which will be deformed into a half-sphere of the same radius $R$ (Figure S1). We assume that the deformation is localized with this disc, so that the change in free energy is given by the change in free energy of this disc. We therefore find the deformation energy to be the difference in the free energy of a disc of radius $R$ and a half-sphere of radius $R$.

The contribution due to tension is given by

$$\Delta F_\sigma = \sigma \Delta A = \sigma \left(2\pi R^2 - \pi R^2\right) = \pi R^2 \sigma \tag{S3}$$

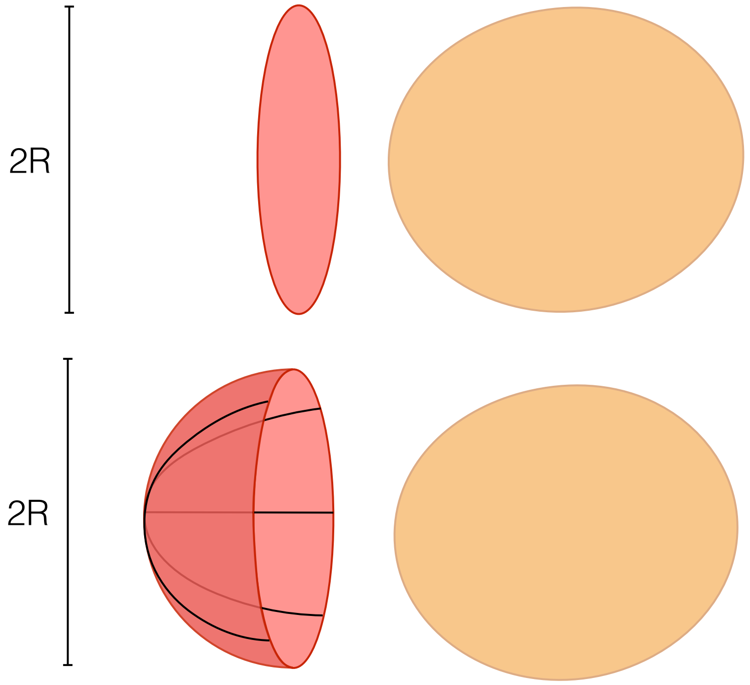


Figure S1: A schematic of the deformation of the patch (pink oval) of erythrocyte membrane as it wraps around the merozoite. In the top half of the figure, the patch—shown in perspective—consists of a disc of radius $R$, where $R$ is also the dimension of the merozoite perpendicular to its axis of symmetry. Following the partial wrapping of the membrane around the merozoite, the membrane is modelled as a half-sphere of the same radius.

Likewise, the contribution due to bending can be written as

$$\Delta F_\kappa \approx 2\kappa \left(2\pi R^2 \frac{1}{R^2}\right) = 4\pi\kappa \tag{S4}$$

where we have omitted the equilibrium bending energy as it is comparatively small.

This yields the result

$$\Delta F = \Delta F_\sigma + \Delta F_\kappa = \pi R^2 \sigma + 4\pi\kappa \tag{S5}$$

$\Delta F$ gives the minimum work needed to wrap the membrane around the merozoite. The receptor-ligand binding energy between erythrocyte and merozoite is likely to contribute energetically [4].

### S1.1 Meaning of bending modulus and tension

The cell membrane has a finite bending modulus, $\kappa$, due to its lipid bilayer structure. This bending modulus corresponds to one's intuition of bending modulus: When one rolls up a sheet of paper, there is no stretching, but doing so requires a minimum amount of work.

In contrast, the tension does not correspond to the naive and intuitive definition of tension. Such a definition of tension would correspond to the energy needed to stretch a flat sheet of membrane, increasing its area. Much like the sheet of paper, the tension of membranes is so high that, in the biological regime, it is equivalent to assuming that the membrane area is fixed, or that the tension diverges.

The measured tension $\sigma$, or "effective tension", is a consequence of the cell having fixed and *excess* surface area. Stretching, therefore, does not increase the intra-lipid distances, but rather suppresses thermal fluctuations [2]. In analogy to the piece of paper, $\sigma$ does not correspond to the enormous tension that would be required to stretch a piece of a paper (and would in any case likely rip it before stretching), but rather the much more manageable tension necessary to stretch a crumpled piece of paper into a smoother piece of paper.
