## Supplementary Information for "The erythrocyte membrane properties of beta thalassaemia heterozygotes and their consequences for *Plasmodium falciparum* invasion"

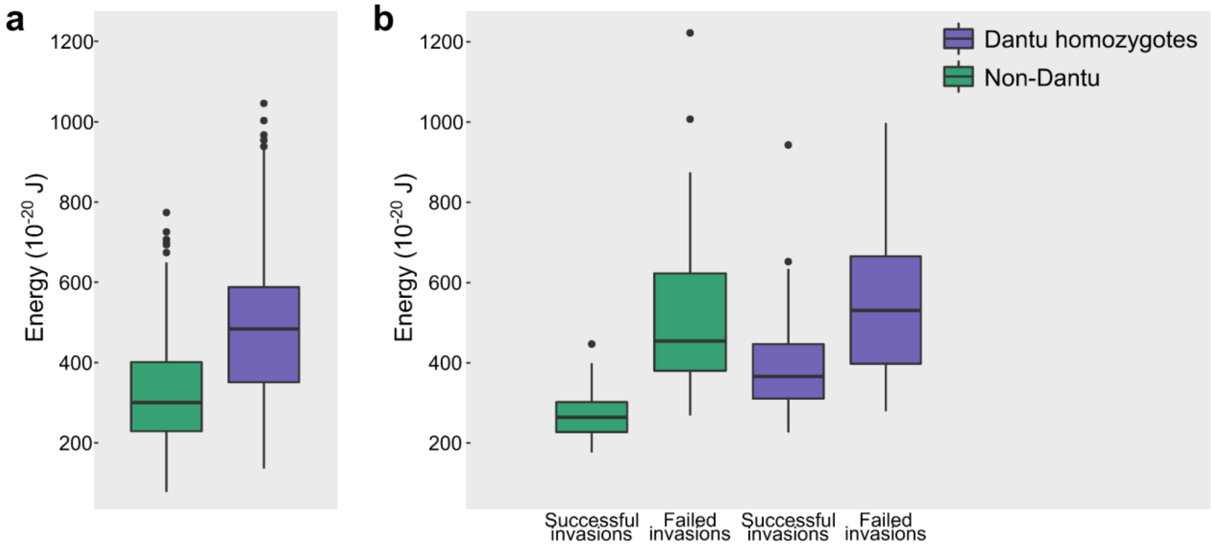


**Figure S1. Wrapping energy of non-Dantu and Dantu homozygotes, for successful and failed invasions.** Panel **a** displays wrapping energy for erythrocytes from Dantu homozygotes and non-Dantu individuals; each boxplot contains data from 5 different people. A mixed model accounting for individual as a random effect and genotype as a fixed effect (see Methods) finds that genotype explains a significant proportion of the variation in wrapping energy (p= 0.0000279), and that the increase in wrapping energy associated with Dantu homozygosity is 151 x $10^{-20}$ J (95% CI 105 x $10^{-20}$ to 202 x $10^{-20}$ J). Panel **b** shows wrapping energy for erythrocytes from non-Dantu individuals and Dantu homozygotes in the case of both successful and failed merozoite invasions. Erythrocytes came from 3 individuals per genotype group. Bending modulus and tension data used to calculate the energy are from Kariuki *et al.*[24], Fig. 3a and 3c. A mixed model accounting for individual as a random effect and genotype and invasion success as fixed effects (see Methods) finds that invasion success explains a significant proportion of the variation in wrapping energy among erythrocytes ($p<2.2 \times 10^{-16}$), and that the difference in wrapping energy associated with failed invasion is an increase of 258 x $10^{-20}$ J (95% CI 210 x $10^{-20}$ to 307 x $10^{-20}$ J).

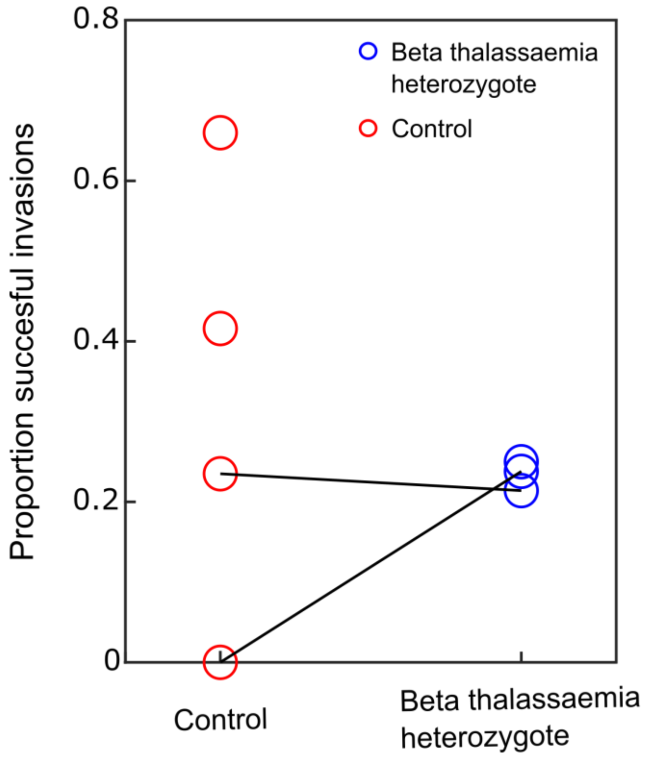


**Figure S2. Proportion of successful invasions by strain 3D7, measured by live imaging video.** The proportion of successful invasion events for 2 matched pairs of beta thalassaemia heterozygote (blue) and control (red) 2018 samples are displayed. We also obtained results for two additional 2018 control samples and one additional 2018 heterozygote sample but not their matched pairs; these are also displayed. Solid lines join heterozygote/control pairs. The number of contact events per sample used to calculate these proportions of successful invasions ranged from 12 to 37, and detailed information on live imaging methods can be found in Methods and in previous publication[24].

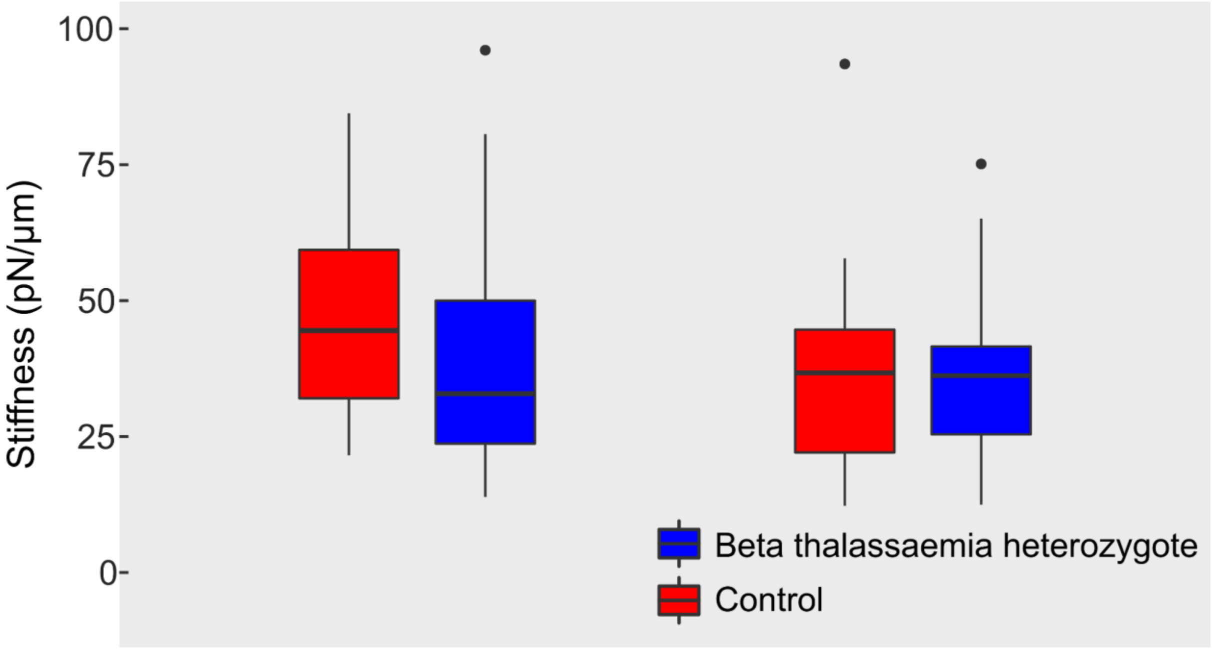


**Figure S3. Erythrocyte stiffness in control and beta thalassaemia heterozygotes.** The boxplots show the erythrocyte stiffness measured at a rate of 1 $s^{-1}$ (see methods) for two pairs of 2019 samples. We find no difference in the stiffness of the control and beta thalassaemia heterozygous cells. These results agree with previously measured stiffnesses for erythrocytes at a strain rate of 1 $s^{-1}$ (Yoon *et al.* [29]) N = 20, 17, 13, 22; Welch t-test p values = 0.36 and 0.89, respectively.
